# CHIR99021 causes inactivation of Tyrosine Hydroxylase and depletion of dopamine in rat brain striatum

**DOI:** 10.1101/2023.05.15.540370

**Authors:** Sally Hamdon, Pol Fernandez-Gonzalez, Muhammad Yusof Omar, Marta González-Sepúlveda, Jordi Ortiz, Carles Gil

**Author notes:** Contributed equally (Jordi Ortiz, Carles Gil). To whom correspondence should be addressed: Carles Gil, Department of Biochemistry and Molecular Biology, School of Medicine, and Institut de Neurociències, Universitat Autònoma de Barcelona, 08193 Bellaterra (Cerdanyola del Vallès, Catalonia), Spain.

## Abstract

CHIR99021, also known as laduviglusib or CT99021, is a Glycogen-synthase kinase 3β (GSK3β) inhibitor, which has been reported as a promising drug for cardiomyocyte regeneration or treatment of sensorial hearing loss. Since the activation of dopamine (DA) receptors regulates dopamine synthesis and they can signal through the β-arrestin pathway and GSK3β, we decided to check the effect of GSK3β inhibitors (CHIR99021, SB216763 and lithium ion) in the control of DA synthesis. Using *ex vivo* experiments with minces from rat brain striatum, we observed that CHIR99021, but not SB216763 nor lithium, causes a complete abrogation of DA synthesis and accumulation at low µM concentrations, pointing to off-target effects of CHIR99021. This decrease can be attributed to tyrosine hydroxylase (TH) inhibition since the accumulation of L-DOPA in the presence of a DOPA decarboxylase inhibitor was similarly decreased. On the other hand, CHIR99021 caused a dramatic increase in the DOPAC / DA ratio, an indicator of DA metabolization, and hindered DA incorporation into striatum tissue to the same extent as tetrabenazine, thus pointing to some effect on DA storage that triggers DA feedback inhibition of TH. In addition, CHIR99021 or SB216763, but not lithium, decreased TH phosphorylation in Ser19, but not in Ser31 or Ser40. These results demonstrate that CHIR99021 can lead to TH inactivation and DA depletion in brain striatum, opening the possibility of its use in DA-related disorders, and shows effects to be considered in future clinical trials.

## 1. INTRODUCTION

Dopamine (DA) transmission is described as essential in the control of behavior through natural rewards, such as food, sex, and nurture (Spanagel and Weiss, 1999). Moreover, dysregulation of DA metabolism is a key event in neurodegenerative diseases, such as Parkinson’s disease (PD) (Johnson et al., 2018) or in mental disorders, such as schizophrenia (Dahoun et al., 2017), addiction (Salamone et al., 2016) or depression (Duda et al., 2020). Moreover, due to the role of DA in the control of coordinated movements, alterations in the dopaminergic system also contribute to diseases causing locomotion impairment (Smith-Dijak et al., 2019). DA is synthetized by tyrosine hydroxylase (TH; tyrosine 3-monooxygenase; E.C. 1.14.16.2) and DOPA decarboxylase and it is rapidly internalized into synaptic vesicles by means of the vesicular monoamine transporter 2 (VMAT2) to prevent its auto-oxidation (Gonzalez-Sepulveda et al., 2023) and to prepare synaptic release (Meiser et al., 2013). Cytosolic DA, including either recently synthetized or reuptaken, can also be metabolized to different compounds including 3,4-dihydroxyphenylacetic acid (DOPAC). Therefore, the ratio between DOPAC and DA levels is usually considered an index of DA metabolism. As an imbalance of brain DA levels could lead to numerous diseases, TH is subjected to an intense regulation. Short-term modulation of TH activity includes end-product feedback inhibition by DA (González-Sepúlveda et al., 2022) and phosphorylation at serine residues Ser8, Ser19, Ser31 and Ser40 (Dunkley et al., 2019). Several studies showed that increased phosphorylation at Ser40 positively correlated with TH activity (Bobrovskaya et al. 2007), while phosphorylation of Ser19 and Ser31 increases the rate of phosphorylation of Ser40 and potentiates TH activation (Lehmann et al., 2006). DA exerts its functions through membrane receptors coupled to G proteins (Thal et al., 2018). DA receptors comprise two classes: the D1-like class (D1 and D5) and the D2-like class (D2, D3 and D4) (Xin et al., 2019). Both classes of receptors can act by two pathways, one inhibits intracellular cAMP via protein G_i_ (Bonifazi et al., 2019) while the other can initiate a cAMP-independent pathway by promoting a scaffolding complex that leads to the activation of both ERK1/2 and GSK3β signaling, through the recruitment of β-arrestin (Jiang et al., 2022). These two pathways are differently turned-on depending on the ligand, in a phenomenon known as biased agonism (Beaulieu & Gainetdinov, 2011).

CHIR99021 (also known as laduviglusib or CT99021) is a GSK3β inhibitor which is currently in clinical phase 2b for the restoration of some types of hearing loss (McLean et al., 2021; ClinicalTrials.gov Identifier: NCT05086276) and is a promising molecule that has also been successfully tested in inhibition of adipogenesis (Bennett et al., 2002), coronary and myocardial diseases (Drakhlis et al., 2021; Badimon et al., 2019), cancer (Oh et al., 2017; Lee et al., 2018; Houben et al., 2022), lung repair therapy (Uhl et al., 2015), stimulation of stem cells (Zhao et al., 2014), and type II diabetes mellitus (Ring et al., 2003). Moreover, CHIR99021 has also been recently proposed as a potential drug for Huntington’s disease (Hu et al., 2021).

In the present work we used *ex vivo* assays with rat brain striatum tissue to study the effects of three GSK3β inhibitors (CHIR99021, SB216763 and lithium) on DA metabolism. The obtained results demonstrate that CHIR99021 indirectly inhibits TH and depletes DA in brain striatum samples and should be considered as a potential drug for the treatment of disorders caused by overproduction of DA. The exact mechanism by which CHIR99021 impairs TH activity is still unknown.

## 2. Materials and methods

### 2.1. Animals

Experiments were conducted with male Sprague-Dawley rats weighing 200-250 g (from the Animal Service, Universitat Autònoma de Barcelona or from Charles River). Animals were housed five per cage with *ad libitum* access to food and water during a 12-hour light / dark cycle. Protocols for animal use were approved by the Ethics Committee for Human and Animal Research (Universitat Autònoma de Barcelona) in accordance with guidelines established by the Ethical Committee for the use of Laboratory Animals in Spain (53/2013) and the European Ethical Committee (2010/63/EU).

### 2.2. Preparation of striatal minces

Freshly obtained rat brains were chilled immediately in modified Krebs-Ringer-bicarbonate medium with the following composition: 120 mM NaCl, 0.8 mM KCl, 2.6 mM CaCl_2_, 0.67 mM MgSO_4_, 1.2 mM KH_2_PO_4_, 27.5 mM NaHCO_3_, and 10 mM glucose, saturated with 95% O_2_ / 5% CO_2_ and adjusted to pH 7.4. Dorsal / medial striata from both hemispheres were dissected and sliced in a 4°C room using a McIlwain tissue chopper obtaining tissue minces of 0.3 x 0.3 mm / side. Tissue minces were suspended in ice-cold Krebs Ringer bicarbonate medium, and washed three times by centrifugation (1,000 g, 1 min, 4°C), to remove debris of damaged cells, and resuspended. Striatal tissue from a single rat yielded up to 30 samples of 25 µL each of the settled minces suspension corresponding to 24 incubation samples and 6 blank samples (0.3 - 0.7 mg protein each). Typically, samples were distributed into 2 mL polypropylene tubes containing 225 µL of ice-cold Krebs Ringer bicarbonate medium. Blank tubes were kept on ice and the rest were incubated at 37°C in soft agitation in an Eppendorf Thermomixer (5 Prime, Inc., Boulder, CO) under 95% O_2_ / 5% CO_2_ atmosphere.

### 2.3. Determination of endogenous L-DOPA, DA and DOPAC levels by HPLC-EC

Levels of L-DOPA, DA and DOPAC were determined in striatal minces. After different incubation times, tissue minces were sonicated in 0.25 M perchloric acid containing 0.25 mM EDTA and 0.1 mM sodium metabisulphite. A 10 µL aliquot was taken for protein quantification by the BCA method (Thermo Fisher Scientific). Samples were spun in an Eppendorf microcentrifuge at for 10 min, and 20 µL of supernatant were used for quantification. The accumulation of DA in the absence of the decarboxylase inhibitor NSD-1015 and of L-DOPA in the presence of 100 µM NSD-1015 was quantified by HPLC with coulometric detection (HPLC-EC) as previously described (González-Sepúlveda et al., 2022). The chromatography system consisted of a reverse-phase C18 column (2.5 µm particle Fortis C18, 10 x 0.46 cm, Sugelabor, Spain) and an ion-pair mobile phase, made up of 100 mM sodium phosphate buffer, 1 mM EDTA, 5 mM octanesulfonic acid (pH 2.5) plus 1 % (v/v) methanol. The flow rate was 1 mL / min. Concentrations of L-DOPA, DA and DOPAC were detected with a Coulochem II (ESA) detector with a model 5011 dual-electrode analytical cell with porous graphite electrodes. The potential of electrodes 1 and 2 was set at -0.05 V and +0.4 V respectively. Standards of L-DOPA, DA and DOPAC at different concentrations (0.05 - 5 pmol / µL) were injected in every experiment to quantify the three metabolites by the external standard method. The obtained concentrations were corrected by protein content in each sample.

### 2.4. Newly-synthesized [_3_H]-DA determination by HPLC-UV

[_3_H]-Tyrosine was purified as described previously (González-Sepúlveda et al., 2022). Concentration-response curve of CHIR99021 on striatal [_3_H]-DA synthesis in minces *ex vivo* was measured after a preincubation period (as in *Preparation of striatal minces* section). CHIR was added using concentrations of 0.3, 3 and 30 μM (n = 6 per concentration) or control without CHIR (n = 6) 10 minutes before the end of preincubation. Then, 0.12 µM purified [_3_H]-tyrosine was added and incubated for 10 additional minutes. In all experiments, [_3_H]-DA synthesis was stopped by the addition of 25 µL of a deproteinizing mixture containing trichloroacetic acid (0.5 % w/v), 1 mM ascorbic acid and 25 nmol DA (internal standard). Samples were homogenized in a Dynatech / Sonic Dismembrator (Dynatech Labs, Chantilly, VA). A 10 µL aliquot was taken for protein quantification by the BCA method (Thermo Fisher Scientific). Tissue homogenates were then centrifuged (12,000 g, 10 min, 4°C), and all supernatants were processed for [_3_H]-DA purification by HPLC-UV. [_3_H]-DA formed during the incubation reaction was separated from [_3_H]-tyrosine by HPLC purification by a modification of previous procedures (Ortiz et al, 2000; Ma et al., 2014). The chromatography system consisted of a reverse-phase C18 column (Tracer Extrasil ODS2, 5 µm particle size, 25 x 0.46 cm; Teknokroma, Spain) and an ion-pair mobile phase, made up of 100 mM sodium phosphate buffer, 1 mM EDTA, 0.75 mM octanesulfonic acid (pH 5) plus 12 % (v/v) methanol. The flow rate was 1 mL / min. This HPLC system completely separates standards of tyrosine and DA detected by UV 285 nm (ring absorbance). Samples contained extremely low levels of radiolabeled tyrosine and DA that were undetectable by UV absorbance. Similarly, endogenous tyrosine and DA were negligible as compared to the amounts of internal standard DA used. The recovery of the internal standard in each sample (internal / external standard peak area) was quantified from internal standard DA HPLC-UV peak areas. 2 mL fractions corresponding to the DA peak were recovered in scintillation vials, mixed with 6 mL Optiphase HiSafe III cocktail, and quantified in a liquid scintillation counter (Perkin Elmer Tri-Carb 2810TR, USA) to determine [_3_H]-DA. Disintegrations per minute (dpm) obtained in HPLC-purified [_3_H]-DA fractions were corrected by DA internal standard recovery and dpm in blank samples. Rate of [_3_H]-DA synthesis was estimated as the ratio of corrected dpm divided by protein content in each incubate and the incubation time in the presence of [_3_H]-tyrosine. Results were expressed as a percentage with respect to control samples run in the same experiment.

### 2.5. Tyrosine Hydroxylase activity in homogenates obtained from striatum

Striata from a single rat were extracted and placed in 5 mL of cold sodium phosphate buffer 10 mM pH 7.4 and homogenized using a glass Potter homogenizer. 200 µL of homogenates were distributed in incubation tubes in the absence or presence of increasing concentrations of CHIR99021. 100 µM NSD-1015, 30 µM tyrosine and 100 µM tetrahydrobiopterin (BH_4_) were added. After 30 min of incubation at 37°C samples were placed in an ice block and 25 µL of a deproteinizing mixture (containing 0.5% w/v trichloroacetic acid) were added. Samples were centrifuged at 10,000 g for 10 minutes and supernatants injected into HPLC-ECD to quantify L-DOPA and expressed as L-DOPA pmol / mg protein.

### 2.6. Western blot

After incubation of striatal minces as described above, Krebs-Ringer buffer was removed by centrifugation and tissue was homogenized in 1% SDS. Protein amount was determined by BCA. Equal amounts of protein were separated by SDS-PAGE electrophoresis followed by transference in polyvinylidene fluoride membrane in a Trans-Blot Turbo Transfer System (Bio-Rad). Blotting buffer contained 25 mM Tris, 200 mM glycine and 10% methanol (v/v). Membranes were blocked for 1 h with Tris-buffered saline, supplemented with 0.1% Tween 20 and 5% (w/v) defatted powdered milk. Then, the membranes were incubated overnight with the indicated antibody diluted in blocking buffer. The primary antibody from rabbit against β-catenin with triple phosphorylation in the residues Ser33, Ser37 and Thr41 was from Invitrogen (PA5-67518, used at 1:1,000), while the mouse antibody against total β-catenin was from Transduction Laboratories (BD610154, used at 1:1,000). The primary antibodies against TH (AB5280, used at 1:2,500), TH phosphorylated at serine 31 (AB5423, used at 1:1,000) and TH phosphorylated at serine 40 (AB5935, used at 1:1,000) were obtained from Millipore. The antibody against TH phosphorylated at serine 19 (AB5935, used at 1:1,000) was obtained from Thermo Fisher Scientific. The secondary antibodies were IRDye 800CW donkey anti-mouse and IRDye 680RD donkey anti-rabbit (Li-Cor) and were used at 1:10,000 in blocking buffer supplemented with 0.01% SDS, for 1 h at room temperature in the dark. Infra-red signals were visualized using an Odyssey Fc Infrared Imaging System (Li-Cor). Total and phosphorylated forms were evaluated on the same membrane and the signals were acquired and quantified with Image Studio Lite software. The ratios of phosphorylated to total protein were calculated respect to controls, which were arbitrarily set to 100%.

### 2.7 Statistical Analysis

Graphs represent incubation samples that may be obtained from different animals, which is important due to the limited amount of brain striatal tissue per animal and its inherent heterogeneity. The use of the term “replicates” has been avoided to stress the heterogeneity of tissue minces randomly assigned to each treatment group. Statistical analysis was carried out with GraphPad Prism9 software (GraphPad Software Inc, USA). Two-way ANOVA was used to analyze the interaction between different factors on synthesis rate (e.g., time, treatment, or concentration of DA as factors). Statistical significance of differences vs. control group was assessed by one-way ANOVA followed by Dunnett’s post-hoc test for dose-response graphs and two-way ANOVA followed by Bonferroni’s post-hoc test for time-response graphs. Sigmoidal dose-response curve regression was used in every case to create the adjusted curves. One-way ANOVA analysis with Newman-Keuls post-test was used in the rest of the experiments.

## 3. Results

### 3.1. CHIR99021 strongly decreases dopamine accumulation and TH activity in rat striatal minces

With the aim of analyzing the effect of CHIR99021 on DA accumulation, striatal minces were treated with 3 different concentrations (0.3, 3 and 30 µM) over an incubation period of 2 hours. A concentration-dependent decay in DA accumulation, with an IC_50_ of 3.76 µM, can be clearly observed (Figure 1A). In fact, the 30 µM concentration was even able to decrease accumulation to the basal level (dashed line), which represents non-incubated tissue kept on ice. 3 µM CHIR99021 also demonstrated a time-dependent effect on DA accumulation (Figure 1B). To further investigate DA metabolism, we also determined the levels of DOPAC and the DOPAC / DA ratio in the presence of growing concentrations of CHIR99021, as DOPAC / DA ratio is considered to reflect the degree of DA metabolization by monoamine oxidase (MAO). While no significant changes in DOPAC levels were found (Figure 1C), a dramatic increase in the DOPAC / DA ratio was detected at 30 µM CHIR99021 (Figure 1D), due to DA decrease. This suggests either an increased DA metabolism or a decreased TH activity. Thus, to assess the effect of CHIR99021 on TH activity, striatal minces or homogenates were incubated in the presence of 100 µM NSD-1015 to inhibit L-aromatic amino acid decarboxylase allowing the quantification of L-DOPA accumulation. While CHIR99021 induced a concentration-dependent inhibition of TH activity in striatal minces with an IC_50_ of 8.16 µM (Figure 1E), no effect of CHIR99021 on TH was observed in striatal homogenates (Figure 1F). These results suggest that the decrease of L-DOPA levels is not a result from direct interaction of CHIR99021 with TH and therefore the intact intracellular machinery is required.

**Figure 1.**
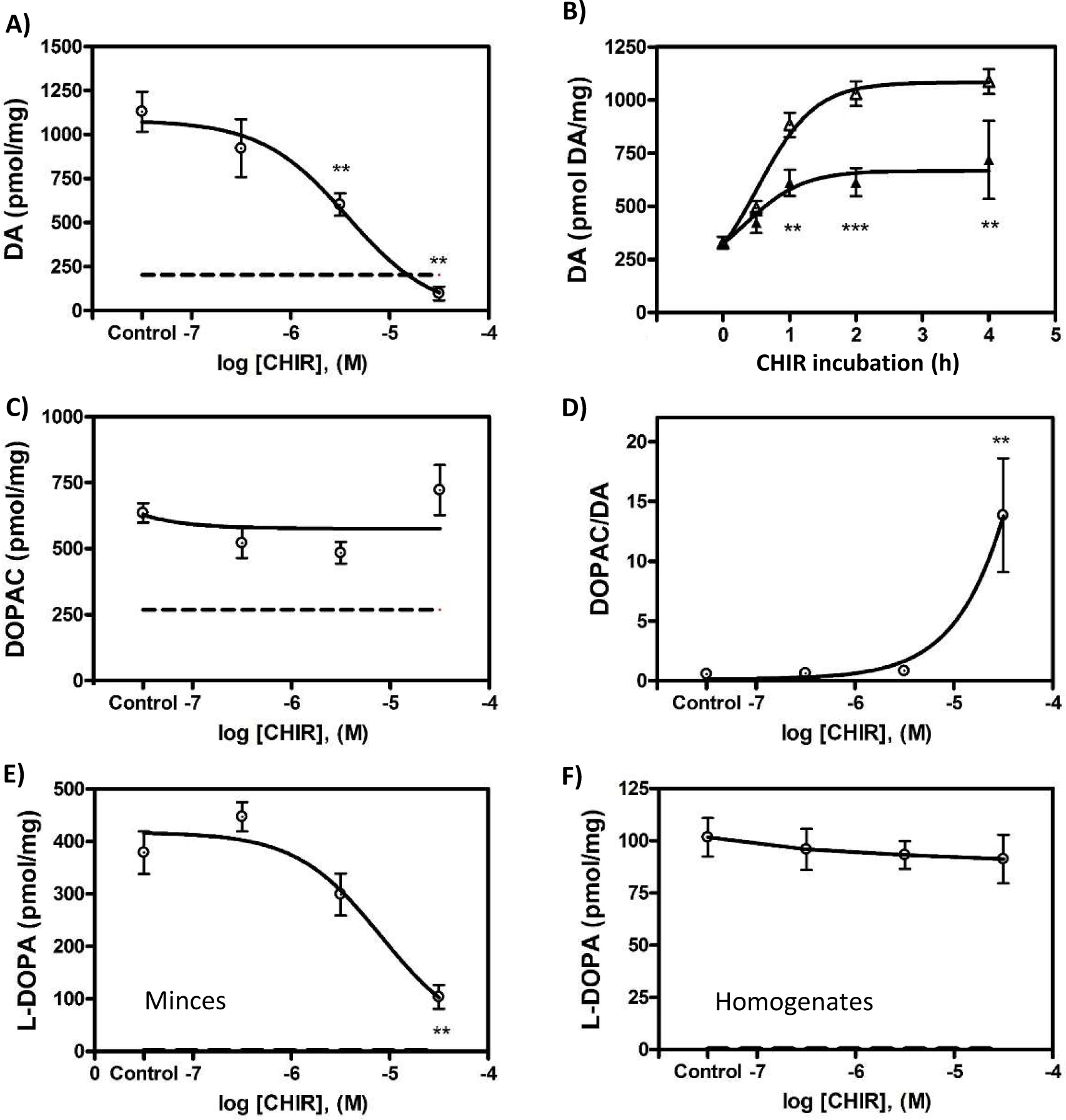
Effects of CHIR99021 on dopamine accumulation and metabolism in rat brain striatal minces. **(A)** Concentration-response curve of CHIR99021 in striatal minces *ex vivo*, measuring dopamine (DA) accumulation over a 2-hour incubation period, using concentrations of 0.3, 3 and 30 μM CHIR99021 or control without CHIR99021 (6 incubates per group obtained from a single animal, n = 6). Basal DA was measured as control samples placed on ice during the 2-hour incubation period (6 incubates, dashed line). **(B)** Time-response curve of DA accumulation in striatal minces *ex vivo* with 3 μM CHIR99021 for 30 minutes, 1 hour, 2 hours, 4 hours (black triangles) and controls (white triangles) for each incubation period. Controls were treated with the corresponding concentration of DMSO used to dissolve CHIR99021 (n = 6), and basal dopamine measured same as in A) and represented as 0 time (n = 6). The same protocol as in A was used in striatal minces to determine DOPAC **(C).** The DOPAC / DA ratio was calculated **(D).** The L-DOPA levels after CHIR99021 treatment were measured in minces **(E)** and in homogenates **(F)** from striatal minces. In the L-DOPA determinations, 100 µM NSD-1015 was also added at the beginning of the experiment. Data are expressed as mean ± SEM for each group, CHIR99021 *vs* control, one-way ANOVA followed by Dunnett’s post-hoc test for dose-response graphs and two-way ANOVA followed by Bonferroni’s post-hoc test for time-response graph. **p<0.01, ***p<0.001. Sigmoidal dose-response curve regression was used in every case to create the adjusted curves.

Since DA accumulation is a result of new DA synthesis and storage, we decided to test the effect of CHIR99021 on newly synthesized DA using a radioisotopic method to measure [_3_H]-DA synthesis from [_3_H]-tyrosine in striatal minces. CHIR99021 decreased [_3_H]-DA synthesis concentration-dependently (Figure 2A), as in the case of DA accumulation. In this case, the value of IC_50_ obtained was approximately 0.30 µM. Western blots against TH were performed, to determine whether the decrease in DA synthesis was due to a decrease in the amount of TH protein. This was not altered under any of the CHIR concentrations tested, either in absolute values or in relation to GAPDH (Figure 2B). Thus, the abrogation of the TH activity by CHIR is not due to loss of TH protein but to a regulatory mechanism affecting efficiency of the enzyme.

**Figure 2.**
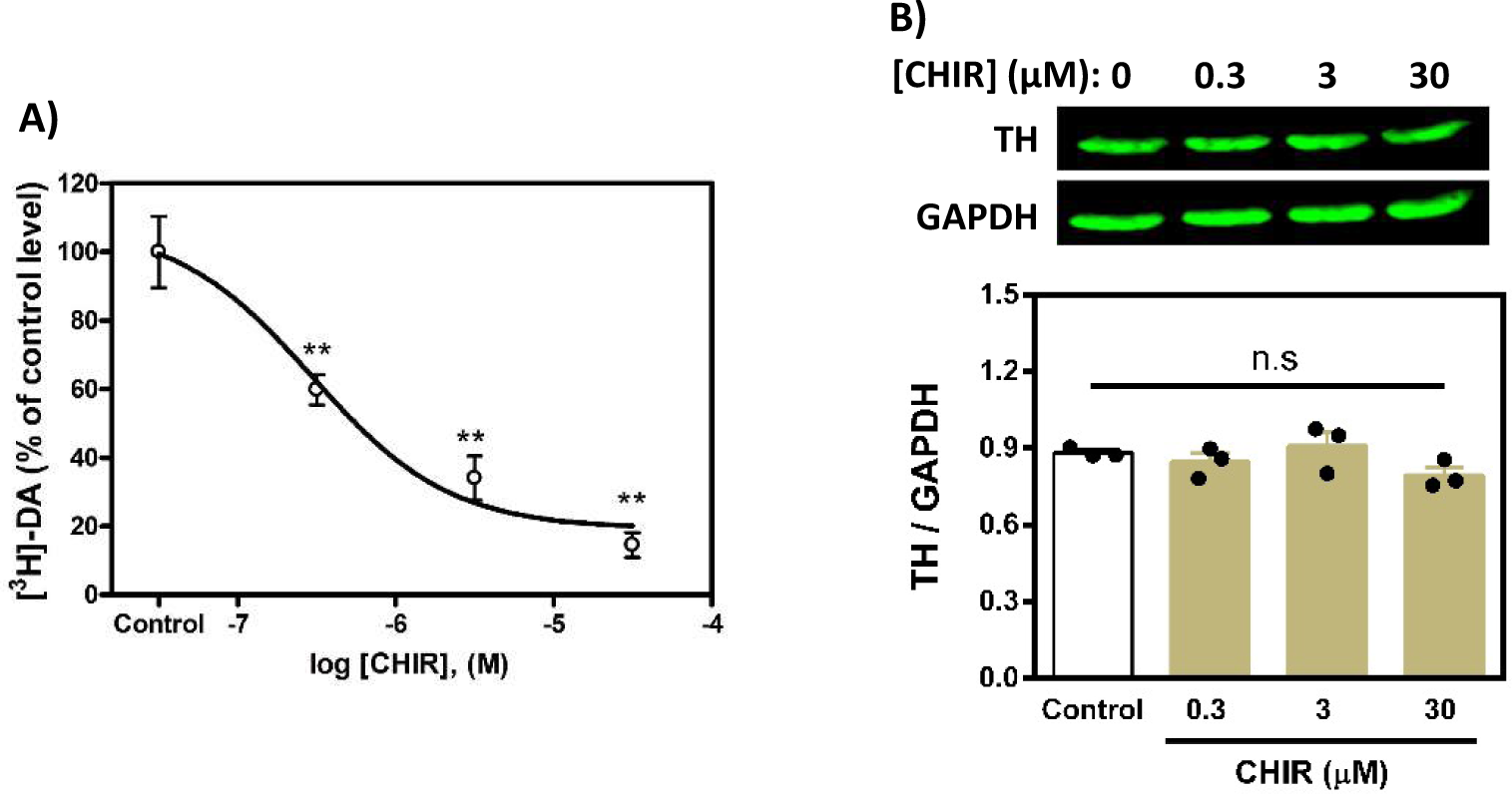
CHIR99021 inhibits tyrosine hydroxylase enzymatic activity. **(A)** Concentration-response curve of CHIR99021 on striatal [_3_H]-DA synthesis in minces *ex vivo* was measured after a preincubation period of 2 hours. CHIR99021 was added using concentrations of 0.3, 3 and 30 μM (n = 6 per concentration) or control without CHIR (n = 6) 10 minutes before the end of preincubation. Then, [_3_H]-tyrosine was added and incubated for 10 additional minutes. The curve is obtained by normalization of the data to percent of control. In all cases, data are expressed as mean ± SEM for each group. Asterisks indicate **p<0.01 CHIR99021 vs control, one-way ANOVA followed by Dunnett’s post-hoc test. The curve was adjusted to sigmoidal dose-response regression. **(B)** TH amount was assessed by western blot after treatment of minces with 0.3, 3 and 30 μM CHIR99021 or control without CHIR99021 (shown as 0), as in A. GAPDH amount was also detected by western blot and TH / GAPDH ratio was calculated and is represented as a bar graph, which shows means and SEM, after One-way ANOVA analysis with Newman-Keuls post-test, which yielded non-significant (n.s.) in every case. Bands from representative western blots are also shown.

### 3.2. SB216763 and lithium ion do not decrease dopamine content or alter DA metabolism

To corroborate a role of GSK3β in the decrease of TH activity and DA accumulation, another GSK3β inhibitor called SB216763 was also tested. SB216763 was able to slightly decrease dopamine accumulation in rat striatal minces over a 2-hour period, but only at 30 µM (Figure 3A) and had no effect at 3 µM or 0.3 µM, even after 4 hours of incubation (Figure 3B). The assessment of TH activity by L-DOPA quantification showed a lack of action by SB216763 at every concentration tested (Figure 3C). Quantification of DOPAC levels at increasing concentrations of SB216763 showed a similar profile as that of DA accumulation shown in 3A, with only a slight decrease of DOPAC amount at 30 µM (Figure 3D), thus yielding a DOPAC / DA ratio unaltered respect to control at any SB216763 concentration (Figure 3E). Given that lithium chloride has antipsychotic properties and has also been reported to inhibit GSK3, the effects of lithium ion on DA metabolism in striatal minces were also tested. We observed that 20 mM LiCl yielded a significant but slight decrease in DA accumulation over a 2-hour incubation period, whereas no effect was detected at 1 or 5 mM (Figure 4A). A time-course with 20 mM lithium showed significant decreases in DA accumulation after 2 and 4 hours of incubation (Figure 4B). Determinations of L-DOPA (Figure 4C), DOPAC (Figure 4D) and DOPAC / DA ratio (Figure 4E) at 1, 5, or 20 mM lithium for 2 hours did not show remarkable changes respect to control.

**Figure 3.**
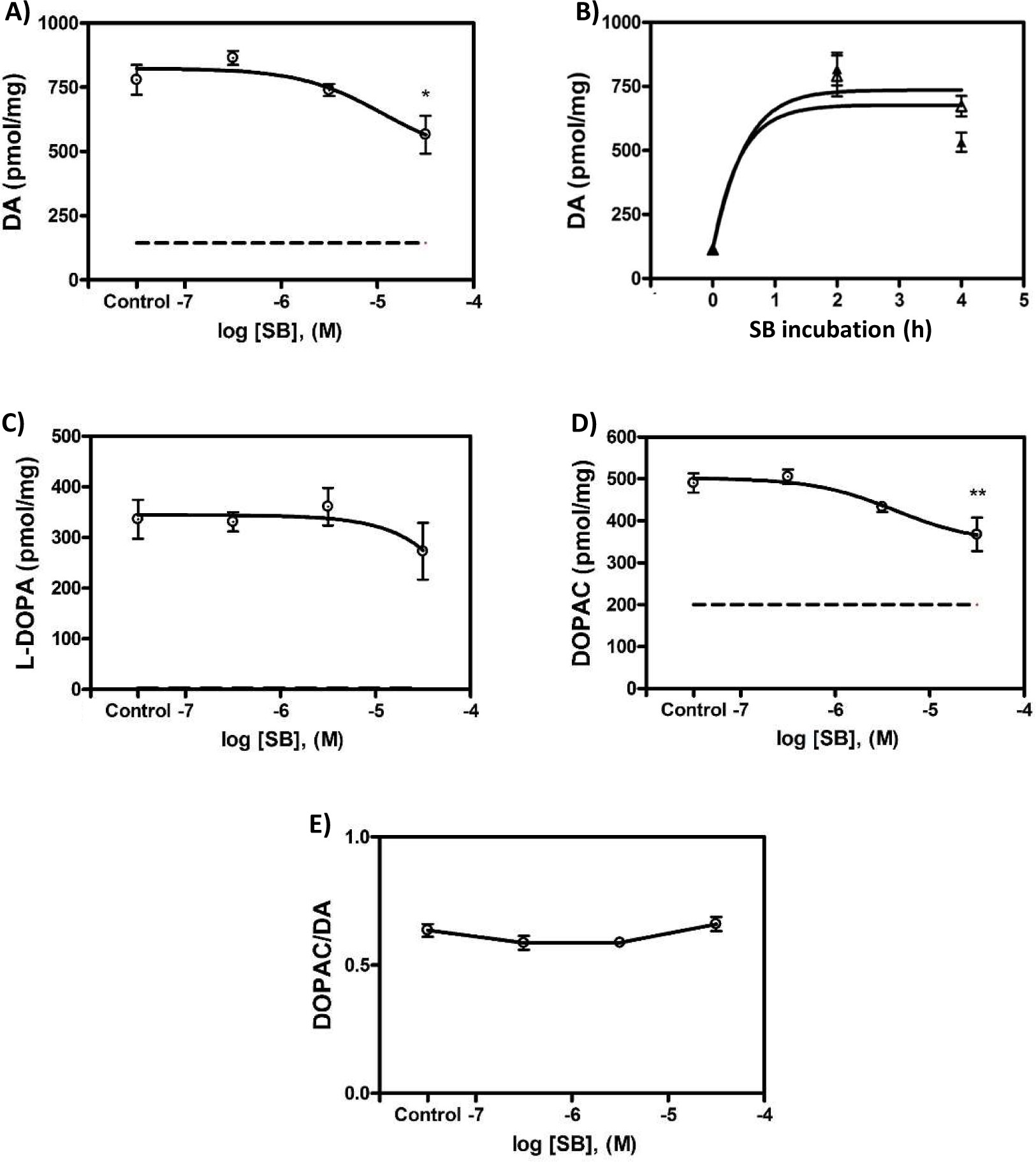
Effect of SB216763 on dopamine accumulation and metabolism in rat brain striatal minces. **(A)** Concentration-response curve of SB216763, measuring dopamine (DA) accumulation over an incubation period of 2 hours *ex vivo*, using concentrations of 0.3, 3 and 30 μM SB216763 or control without SB217663 (6 incubates per group obtained from a single animal, n = 6). Basal dopamine was measured as control samples placed on ice during the 2-hour incubation period (6 incubates, dashed line). **(B)** Time-response curve of DA accumulation on striatal minces *ex vivo* for a SB216763 concentration of 3 μM and for 2 and 4 hours, and controls for each incubation period, which were treated with the corresponding concentration of DMSO used to dissolve SB216763 (n = 6), and basal dopamine measured same as in (A) and represented as 0 time (n = 6). The same protocol as in A was used in striatal minces to determine L-DOPA **(C)** and DOPAC levels **(D)**. The DOPAC / DA ratio was calculated **(E)**. Data are expressed as mean ± SEM for each group, SB216763 vs control, one-way ANOVA followed by Dunnett’s post-hoc test for dose-response graph and two-way ANOVA followed by Bonferroni’s post-hoc test for time-response graph. *p<0.5, **p<0.01. Sigmoidal dose-response curve regression was used in every case to create the adjusted curves.

**Figure 4.**
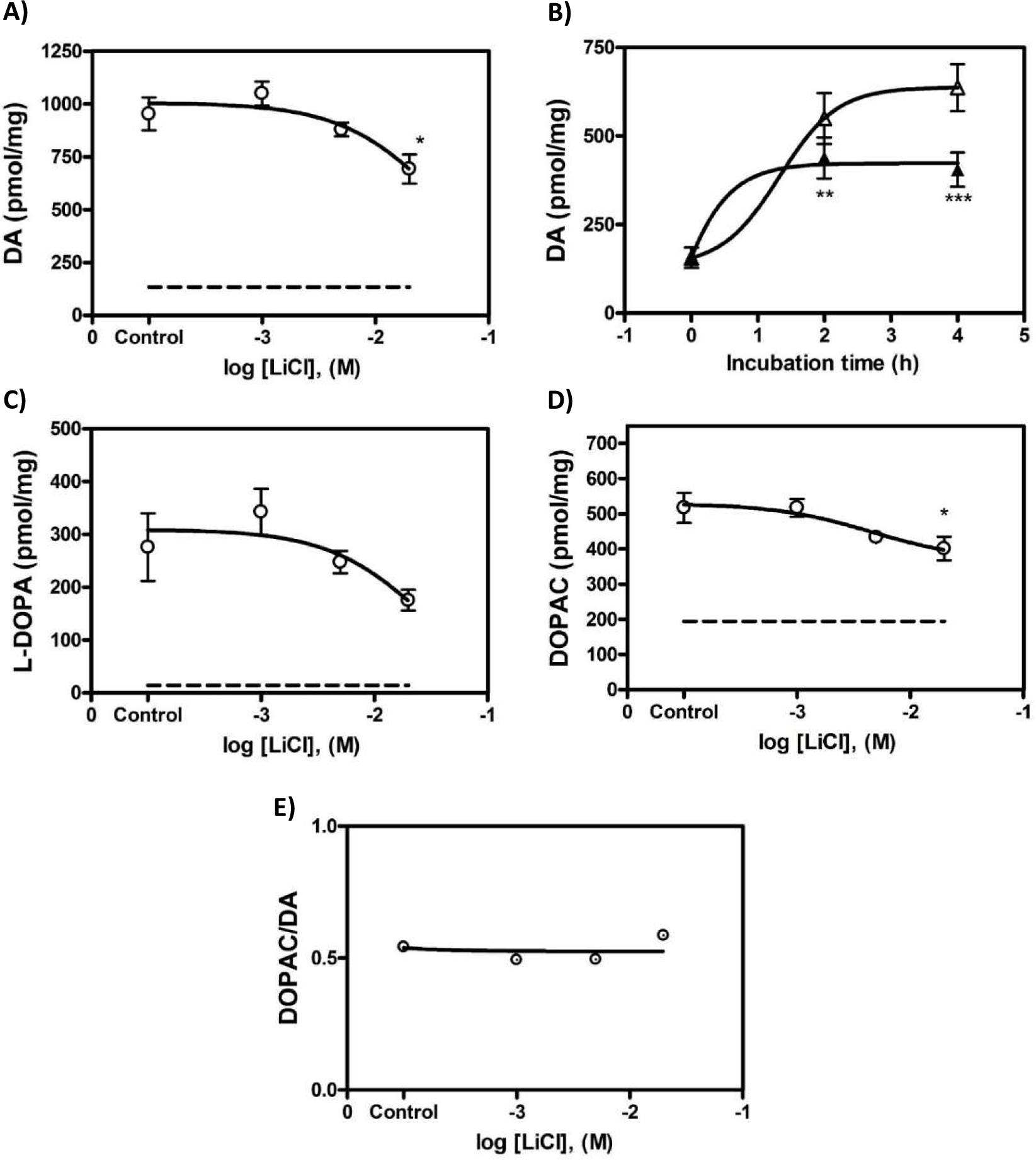
Effect of lithium ion on dopamine accumulation and metabolism in rat brain striatal minces. **(A)** Concentration-response curve of lithium, measuring DA accumulation over an incubation period of 2 hours *ex vivo*, using concentrations of 1, 5 and 20 mM LiCl or control without lithium (6 incubates per group obtained from a single animal, n = 6). Basal DA was measured as control samples placed on ice during the 2-hour incubation period (6 incubates, dashed line). **(B)** Time-response curve of DA accumulation on striatal minces *ex vivo* for a lithium concentration of 20 mM for 2 and 4 hours (n = 6), and basal DA measured same as in (A) and represented as 0 time (n = 6). The same protocol as in A was used in striatal minces to determine L-DOPA **(C)** and DOPAC levels **(D)**. The DOPAC / DA ratio was calculated **(E)**. Data are expressed as mean ± SEM for each group, lithium vs control, one-way ANOVA followed by Dunnett’s post-hoc test for dose-response graph and two-way ANOVA followed by Bonferroni’s post-hoc test for time-response graph. *p<0.5, **p<0.01, ***p<0.001. Sigmoidal dose-response curve regression was used in every case to create the adjusted curves.

### 3.3. CHIR99021 and SB216763, but not lithium, inhibit GSK3β activity on β-catenin in striatum minces

GSK3β phosphorylates β-catenin in the residues S33, S37 and T41. Thus, to directly test the inhibitory effect of CHIR99021, SB216763 and of lithium on the GSK3β activity in our system, the amount of triple phosphorylated β-catenin was assessed by western blot. Near infrared-labeled secondary antibodies were used, thus phosphorylated and total protein levels could be assessed in the same membrane, giving rise to reliable signal quantifications. Treatment with 3 µM CHIR99021 and with 3 µM SB216763 for 2h decreased GSK3β-phosphorylated β-catenin by half approximately, whereas 5 mM lithium for 2h did not show any effect on β-catenin phosphorylation (Figures 5A and 5B). These results corroborate that CHIR99021 and SB216763 cause inhibition of GSK3β in our striatal minces. In consequence, the lack of SB216763 effect on DA metabolism is due to absence of GSK3β action on TH and not to a lack of compound efficacy.

**Figure 5.**
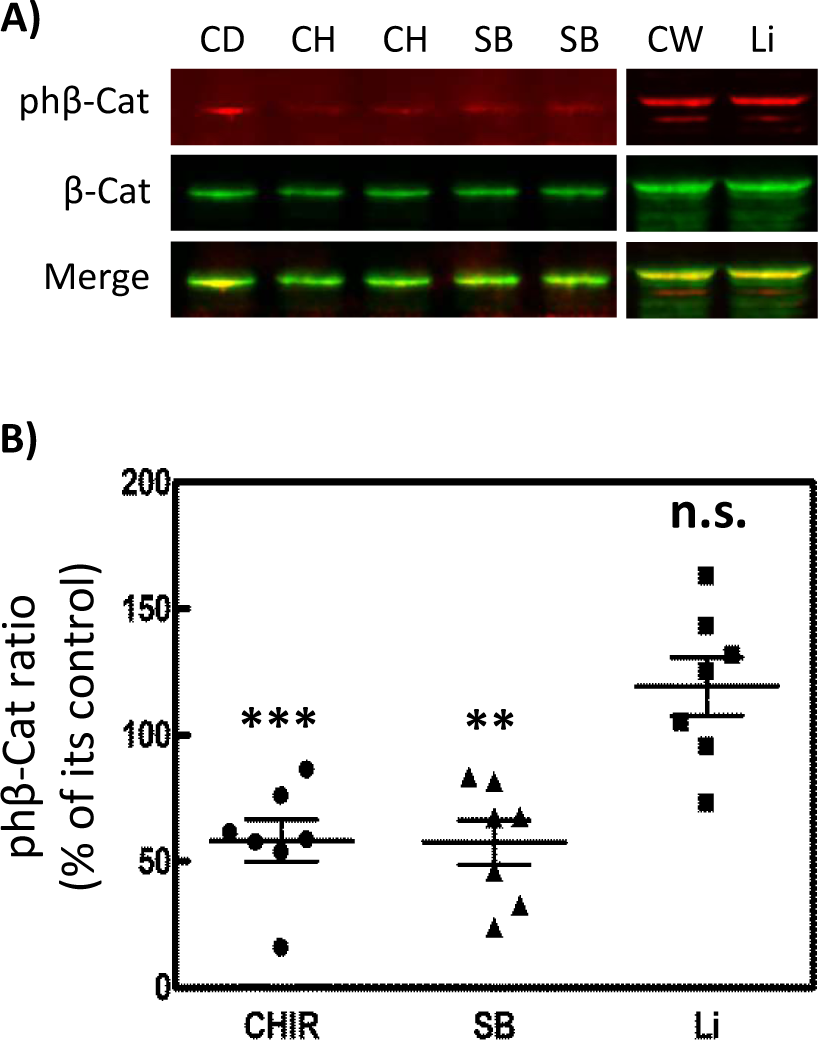
GSK3β inhibitors CHIR99021 and SB216763, but not lithium, cause decrease in phosphorylation of β-catenin in rat striatal minces. Tissue samples were treated with two GSK3 inhibitors, CHIR99021 (CH, 3 μM) or SB216763 (SB, 3 μM), both for 2h. In parallel, striatal minces were also treated with lithium cloride (Li, 5 mM) for 2h. Levels of phosho-β-catenin were assessed with western blot using NearIR-labeled secondary antibodies, as described in Materials and Methods section. **(A)** Representative western blots are shown after CHIR99021, SB216763 or lithium treatment. Red signal corresponds to β-catenin triple phosphorylated in S33, S37 and T41, while green signal corresponds to total β-catenin. Merge of both signals appears in yellow. Control with only DMSO as vehicle was used in the case of CHIR99021 and SB216763 (shown as CD), while control with water as vehicle was used in the case of lithium (shown as CW). **(B)** Scatter plot representation of the β-catenin phosphorylation ratios (phoshorylated over total) after treatment with CHIR99021, SB216763 or lithium. Each point corresponds to the percentage of the ratio respect to its control (CD or CW). Graphics shows means ± SEM, after One-way ANOVA analysis with Newman-Keuls post-test. (** p<0.01, *** p<0.001, n.s. non-significant).

### 3.4. CHIR99021 and SB216763 decrease TH phosphorylation in serine 19, but not in serines 31 and 40

Since phosphorylation in serine residues have been described as a regulatory event of TH activity, we explored the effect of CHIR99021 on TH phosphorylation. Western blots with phosphospecific antibodies showed that GSK3β inhibition cause significant decrease of TH phosphorylation in serine 19, approximately 30% after CHIR99021 or SB216763 treatment for 2h (Figures 6A and A’). On the other hand, no changes in phosphorylation were detected neither in serine 31 nor in serine 40 (Figures 6B, 6B’, 6C and 6C’). Treatment with LiCl (5 mM for 2h) did not yield change in any of the tested phosphosites.

**Figure 6.**
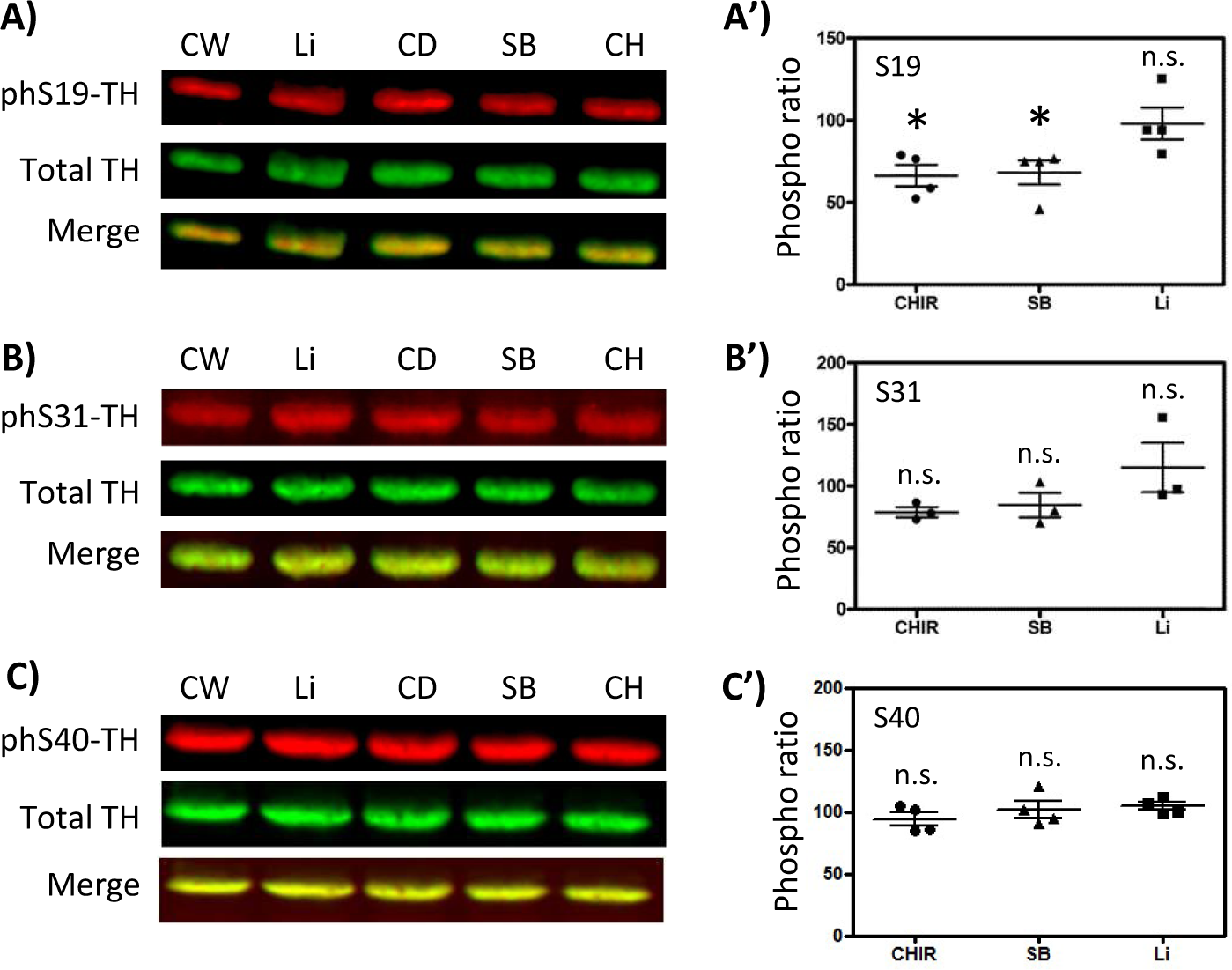
GSK3 inhibitors decrease phosphorylation of Tyrosine Hydroxylase in Ser19, but not in Ser31 or Ser40. Samples from rat striatal minces were treated for 2 hours with CHIR99021 (CH, 3 μM), SB216763 (SB, 3 μM), lithium chloride (Li, 5 mM), DMSO (control for CH99021 and SB216763, shown as CD) and water (control for LiCl, shown as CW). Phosphorylation in every case was assessed by western blot and NIR-labeled secondary antibodies. Red signal corresponds to phosphorylated TH, green signal corresponds to total TH, while merge of both signals appears in yellow. Representative western blots of phosphorylated TH in Ser19 **(A)**, Ser31 **(B)** and Ser40 **(C)**, after treatment with CHIR99021, SB216763 and lithium ion are shown. Next to each group of western blots, the corresponding Scatter plot representations of every phosphorylation ratio (phospho-TH signal over total TH signal) are shown, in percentage respect to its respective control **(A’, B’ and C’)**. Graphics show mean ± SEM, after One-way ANOVA analysis with Newman-Keuls post-test. (*p<0.5, n.s. non-significant).

### 3.5. CHIR99021 impairs the accumulation of exogenous DA into striatal minces

To further explore the mechanisms that lead to the dramatic action of CHIR99021 on DA accumulation, we tested the effect of CHIR99021 on DA transport into striatal tissue. Striatal minces were incubated with 5 µM DA in the absence or presence of CHIR99021 30 µM or of tetrabenazine (TBZ) 1 µM, a VMAT2 inhibitor, and the amount of DA incorporated inside tissue was determined by HPLC, as well as the amount of DOPAC (Figure 7A). Incubation of minces with exogenously added DA caused a large increase in the DA found in the tissue, which is interpreted as DA incorporation into tissue, mainly synaptic vesicles thanks to VMAT2 activity. Pretreatment with either CHIR99021 or TBZ similarly caused a clear reduction (≈ 70%) of the DA additionally incorporated into tissue respect to the control situation (Figure 7B). On the other hand, the assessment of DOPAC levels showed a large increase after DA incubation, but they were poorly or non-modified after TBZ or CHIR99021 pretreatment (Figure 7C). The DOPAC / DA ratios after every treatment were also assessed (Figure 7D), showing that DA turnover is moderately increased by TBZ and CHIR909021 when exogenous DA is present. These results point to an effect of CHIR99021 on some component of the DA transport or storage inside cells.

**Figure 7.**
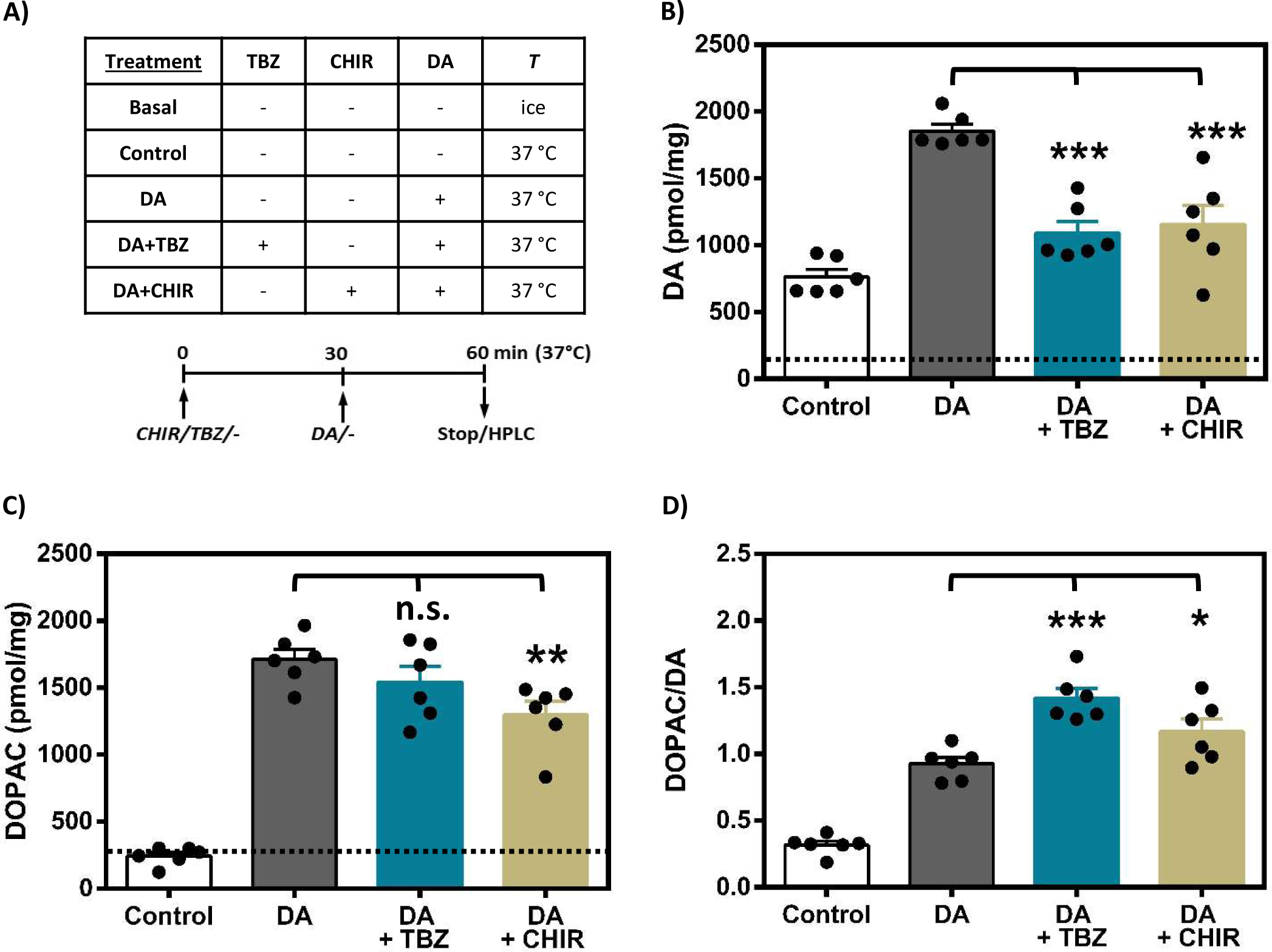
CHIR99021 decreases dopamine storage inside rat brain striatal minces. **(A)** Striatal minces were incubated for 1 hour in ice (Basal) or at 37°C with tetrabenazine 1 µM (TBZ), with CHIR99021 30 µM (CHIR) or with Krebs-Ringer buffer (-). For the last 30 min of incubation 5 µM DA was also added, except in the Basal and Control samples. The amount of DA **(B)** and of DOPAC **(C)** inside tissue was determined at the end of the incubations in every case by HPLC (as pmol DA / mg protein, 6 incubates per group obtained from a single animal). Basal DA or DOPAC in untreated samples is indicated in both cases as dashed lines. Control samples were incubated at 37°C without any drug or DA. The DA turnover was also determined in every case as DOPAC / DA ratios **(D)**. Data are expressed as mean ± SEM for each group. One-way ANOVA test was performed followed by Newman-Keuls post-test vs the DA alone treatment (DA) when indicated (n.s., non-significant, *p<0.5, **p<0.01, ***p<0.001).

## 4. Discussion

CHIR99021 is a small organic molecule that potently inhibits GSK3α (IC_50_∼10 nM) and GSK3β (IC_50_∼5 nM) with high selectivity and is considered the standard activator of the Wnt/β-catenin pathway (Bennett et al., 2002). Moreover, it is a compound that has been tested in a plethora of pathologies (Drakhlis et al., 2021; Badimon et al., 2019; Oh et al., 2017; Lee et al., 2018; Houben et al., 2022; Uhl et al., 2015; Zhao et al., 2014; Ring et al., 2003), showing pharmacological potential in some of them. Furthermore, treatment of isolated rat pancreatic islets with CHIR99021 or SB216763 increases the rate of beta cell replication (Mussmann et al., 2007). Even though CHIR99021 is considered the gold standard for GSK3β inhibition, its off-target effects have been demonstrated. A report describes some action of CHIR99021 on more than 20 kinases as well as the existence of more specific GSK3 inhibitors (An et al., 2012). These off-target effects have been highlighted by a recent report showing that CHIR99021 causes, independently of GSK3, the suppression of the proteasomal degradation of CAST and of the related calpain activity (Hu et al., 2021). This suppression leads to a great enhancement of the mitochondrial function by impairing the action of the GTPase dynamin-related protein 1 (Drp1) on mitochondrial fragmentation. The authors hypothesized that CHIR99021 suppresses CAST degradation through inhibition of ubiquitin ligases or activation of deubiquitinases (Hu et al., 2021). Additionally, the promotion by CHIR99021 of mitochondrial biogenesis, oxidative phosphorylation, and the production of reactive oxygen species in human endodermal progenitor cells has also been observed, in spite that these effects were attributed to the GSK3 / β-catenin pathway and not to off-target effects of CHIR99021 (Ma et al., 2019).

The feedback inhibition of DA synthesis by catecholamines, DA itself but also adrenaline and noradrenaline, has been described (Daubner et al., 2011). Computational analyses have indicated the potential importance of this feedback effect in dopaminergic neurotransmission (Wallace, 2007), while *ex vivo* experimental studies have recently revealed that spontaneous inhibition in DA synthesis appeared when storage reached saturation, causing DA spillover and end-product feedback-inhibition of TH in living tissue (González-Sepúlveda et al., 2022). *In vitro* studies suggest the existence in TH of two DA binding sites with high and low affinities (K_D_ 4 nM and 90 nM, respectively) (Gordon et al., 2008; Nakashima et al., 2009; Dickson and Briggs, 2013; Tekin et al., 2014). The TH low-affinity site / conformation could act as a physiological sensor in case that synaptic vesicles were filled with DA at its maximum capacity, since the surplus of the DA inside the cytosol of synaptic terminals would bind to the low-affinity site / conformation, causing inhibition of TH activity. In agreement with the role of DA in TH inhibition by CHIR99021 is the observation that the determination of the TH activity in the basis of L-DOPA levels under the presence of the decarboxylase inhibitor NSD-1015 (Figure 1E) yielded an IC_50_ one order of magnitude higher than when TH activity was assessed by _3_H-DA production (Figure 2A). Thus, the enabling of DA production potentiates the depleting effect of CHIR99021 in terms of efficacy. Additionally, feedback TH inhibition by other catecholamines cannot be discarded.

Regarding the effect of both CHIR99021 and of SB216763 on TH phosphorylation, Ser19 is reported to be phosphorylated in a 14-3-3 dependent-manner and has little effect on the activity of TH in vitro (Itagaki et al. 1999; Toska et al. 2002). Phosphorylation of TH in Ser19 is related to enzyme degradation of the enzyme by proteasome (Nakashima et al. 2016). The interaction with 14-3-3 proteins, mainly with the γ isoform (Halksau et al., 2009), reduces the sensitivity of phosphorylated human TH isoform 1 to proteolysis by protecting its N-terminal part (Obsilova et al. 2008). Despite the afore mentioned, the dramatic abrogation of TH activity shown in the present work cannot be caused by loss of TH protein, since its levels remain unaltered after CHIR99021 treatment, nor to the relatively small change in phosphoSer19.

One relevant finding of the present work is the dramatic increase of the DOPAC / DA ratio caused by CHIR99021 (Figure 1D). This result can be explained by the enhancement of the MAO activity triggered by the accumulation of its substrate, i.e., DA in the cytosol of the synaptic terminal. This DA accumulation would lead to the inhibition of TH by a feed-back effect, as has been described by our group (González-Sepúlveda et al, 2022) and previously mentioned. An accumulation of DA in striatal minces could also be explained by impairment of the VMAT2 transporter, as it has also been observed with TBZ in a previous report from our group (González-Sepúlveda et al, 2022), giving rise to the possibility that CHIR99021 exerts an off-target inhibitory effect on VMAT2, besides acting as a GSK3β inhibitor. The comparison of the effects of CHIR99021 and TBZ on the transport of extracellular DA inside striatal minces shows a relevant similarity (Figure 7). Since the main portion of the DA in the nerve terminal resides inside synaptic vesicles, where it is protected from metabolism prior to their synaptic release, we can assume that most of the DA incorporated into striatal minces directly reflects the amount of DA inside synaptic vesicles. TBZ is used for the treatment of the motor symptoms in Huntington’s disease (HD), a fatal neurodegenerative disorder, and other hyperkinetic disorders (Jankovic, 2009), but it can cause side effects, like nausea, difficulty in speaking, severe muscle stiffness or, even, depression (Jankovic, 2016). In any case, the reduction of DA release in the motor striatum by VMAT2 inhibition is considered a good therapeutic approach to reduce the involuntary hyperkinetic movements of tardive dyskinesia (Stahl, 2018). In fact, CHIR99021 showed to potently improve mitochondrial function and enhance cell viability in several models of HD (Hu et al., 2021). Although the exact mechanism of CHIR-mediated TH inhibition has not been fully elucidated, its interference with DA storage should be seriously considered. Thus, an available option to TBZ in the hands of the doctors would be a benefit in cases with severe side effects, and CHIR99021 appears as a candidate to be also used in HD or other movement pathologies, such as tardive dyskinesia or Tourette syndrome.

## CRediT authorship contribution statement

**Sally Hamdon:** Methodology, Investigation, Formal analysis, Writing - Original Draft, Visualization. **Pol Fernandez-Gonzalez:** Methodology, Investigation, Formal analysis. **Mohammed Yusof Omar:** Methodology, Investigation, Formal analysis. **Marta González-Sepúlveda:** Visualization, Formal analysis, Writing - Original Draft. **Jordi Ortiz:** Conceptualization, Validation, Formal analysis, Writing - Original Draft, Visualization, Supervision, Project administration, Funding acquisition. **Carles Gil:** Conceptualization, Validation, Formal analysis, Writing - Original Draft, Visualization, Supervision, Project administration, Funding acquisition.

## Declaration of Competing Interest

The authors declare that they have no known competing financial interests or personal relationships that could have appeared to influence the work reported in this paper.

## Abbreviations

DA: dopamine
L-DOPA: levodopa
DOPAC: 3,4-dihydroxyphenylacetic acid
TH: tyrosine hydroxylase
VMAT2: vesicular monoamine transporter 2
TBZ: tetrabenazine
GSK3: Glycogen-synthase kinase

## Acknowledgements

This work was supported by Spanish Government grant SAF2017-87199-R. S.H. received a predoctoral fellowship from the Universitat Autònoma de Barcelona. We thank the skillful technical assistance of Susana Benítez

## Notes

### Competing Interest Statement

The authors have declared no competing interest.

### Summary of Updates

Title has been changes to strength some points; Parts of the Introduction, of the Results and of the Discussion sections have been updated to make them clearer; Figure 1 sections have been reordered; Figure 7 has been improved with new sections and individual data points; A new author has been added.

